# Machine-Learning Prediction of Comorbid Substance Use Disorders in ADHD Youth Using Swedish Registry Data

**DOI:** 10.1101/661983

**Authors:** Yanli Zhang-James, Qi Chen, Ralf Kuja-Halkola, Paul Lichtenstein, Henrik Larsson, Stephen V Faraone

## Abstract

**Background:** Children with attention-deficit/hyperactivity disorder (ADHD) have a high risk for substance use disorders (SUDs). Early identification of at-risk youth would help allocate scarce resources for prevention programs.

**Methods:** Psychiatric and somatic diagnoses, family history of these disorders, measures of socioeconomic distress and information about birth complications were obtained from the national registers in Sweden for 19,787 children with ADHD born between 1989-1993. We trained 1) cross-sectional machine learning models using data available by age 17 to predict SUD diagnosis between ages 18-19; and 2) a longitudinal model to predict new diagnoses at each age.

**Results:** The area under the receiver operating characteristic curve (AUC) was 0.73 and 0.71 for the random forest and multilayer perceptron cross-sectional models. A prior diagnosis of SUD was the most important predictor, accounting for 25% of correct predictions. However, after excluding this predictor, our model still significantly predicted the first-time diagnosis of SUD during age 18-19 with an AUC of 0.67. The average of the AUCs from longitudinal models predicting new diagnoses one, two, five and ten years in the future was 0.63.

**Conclusions:** Significant predictions of at-risk co-morbid SUDs in individuals with ADHD can be achieved using population registry data, even many years prior to the first diagnosis. Longitudinal models can potentially monitor their risks over time. More work is needed to create prediction models based on electronic health records or linked population-registers that are sufficiently accurate for use in the clinic.

## Introduction

In recent years, prevalence of substance use disorders (SUD) have increased significantly (1, 2), magnifying the impact of many adverse consequences (3-9). From 2005 to 2015, death due to opioid, cocaine and amphetamine use disorders increased 30-68% (10). In 2015, over 306,000 deaths were caused by SUD globally, which is 26 times of the total deaths caused by natural disasters and 44% more than all deaths caused by forces of war, violence and legal interventions (10).

Twin studies showed that genes and their interaction with the environment constitute 50-75% of the liability to develop SUD (11, 12). Many risk-modifying environmental factors have been studied including stress and trauma in early life and family, education, socioeconomic status (SES) and cultural influences (12-19). Having attention deficit/hyperactivity disorder (ADHD) is associated with a significantly increased risk for later SUDs (20-22). Relatives of individuals diagnosed with ADHD also have a higher risk for SUDs (23). A large study on the UK Biobank data (N= 135,000) found that the polygenic risk for ADHD significantly predicted alcohol and nicotine use (24). Furthermore, ADHD symptoms, such as inattention, hyperactivity and impulsivity, can cause behavioral problems and stress at home and school, which in turn can increase the risk for substance use.

For individuals with ADHD, however, stimulant therapy was found to decrease the rates of smoking and other SUDs later in life (25, 26) (27, 28). Behavior therapy was also found to significantly reduce substance use reported by ADHD youth at least to 24-month follow-up (The Multimodal Treatment Study of Children with ADHD (MTA) study, (29)). These data suggest that early identification of ADHD youth who are at risk for SUD would allow for more targeted early interventions and possibly prevention of future SUD. Recent studies have demonstrated the feasibility of developing risk prediction models in psychiatry, but the literature is very limited, and most studies are based on small samples (30, 31). Few studies have applied machine-learning algorithms to large-scale data from electronic health records (32) or linked population-registers, and no previous study using such data has been performed in the context of ADHD. Moreover, many prior machine learning studies in psychiatry have been limited by small samples sizes and lack of appropriate replication samples (see, for example, Zhang-James’ et al. review of machine learning applications to neuroimaging data (33).

To address these limitations, the present study sought to develop prediction models using machine learning algorithms to identify ADHD youth at-risk for SUDs. We used information available from the Swedish population registries to construct potential predictors, including the medical history of psychiatric and somatic illnesses for the index children and their immediate family members, as well as their available perinatal records, and socioeconomic, educational and geographic data. Our goal was to determine if the information from the registries could be used to train a clinically useful machine learning algorithm to accurately identify youth with ADHD at risk for SUDs.

## Methods

### Data sources and study population

The study was approved by the Regional Ethical Review Board in Stockholm, Sweden. The sample comprised 19,787 children born between 1989-1993 who had a life-time diagnosis of ADHD (National Patient Registry (NPR) ICD-9: 314; ICD-10: F90). We excluded those: 1) who died or emigrated prior to age 20, 2) who had no father’s information or 3) who had no socioeconomic records during the ages 0 to 18. The final dataset contained 19,184 individuals. SUD was defined by either a diagnosis (ICD-9 304, 305, 306; ICD-10: F10-19) or a prescription of medication for SUD treatment (Supplementary Table 1). We considered the first diagnostic or prescription record as the “onset” of SUD in this study. Supplementary Table 2 shows that 10% of the sample had SUD onset prior to age 17, 8.8% had SUD diagnoses recorded during the ages 18 and 19, and 5.9% of the total sample had a SUD onset during age 18 to 19.

**Table 1.** Top 20 important features.

| Rank | Importance | Features |
| --- | --- | --- |
| 1 | 25% | SUD diagnosis (index child: 12-17) |
| 2 | 10% | Non-Violent Crimes (index child: 12-17) |
| 3 | 6% | Violent Crimes (index child: 12-17) |
| 4 | 3% | ADHD Diagnosis (index child: 2-12) |
| 5 | 2% | Psychostimulants treatment (index child: 12-17) |
| 6 | 2% | Family Income (min percentile: 12-17) |
| 7 | 2% | Anxiety diagnosis (index child: 12-17) |
| 8 | 1% | Non-Violent Crimes (father: prenatal) |
| 9 | 1% | NDEP* (max score:12-17) |
| 10 | 1% | Family Income (max percentile: 12-17) |
| 11 | 1% | Family Income (mean percentile: 12-17) |
| 12 | 1% | Family Income (min percentile: 2-12) |
| 13 | 1% | NDEP (mean score: 0-2) |
| 14 | 1% | Family Income (min percentile: 0-2) |
| 15 | 1% | Family Income (max percentile: 0-2) |
| 16 | 1% | NDEP (max score: 0-2) |
| 17 | 1% | Family Income (mean percentile: 0-2) |
| 18 | 1% | Family Income (mean percentile:2-12) |
| 19 | 1% | ADHD Diagnosis (index child: 12-17) |
| 20 | 1% | NDEP (max score: 2-12) |

\*NDEP, Neighborhood deprivation scores
Note: Importance score represents percentage of contribution toward the prediction accuracy.

### Predictors and missing information

The following registers were used to extract linked data using the unique personal identification number assgined for each Swedish national (34): 1) Medical Birth Register, which was established in 1973 and includes data on prenatal and perinatal measures of all births in Sweden (35); 2) National Patient Register, which contains inpatient care since 1964 (psychiatric care since 1973) and outpatient visits to specialty care since 2001 (36); 3) Total Population Register, which provides information on life events such birth, death, migration and family relationships (37); 4) Multi-Generation Register, which constitutes a part of the Total Population Register, but links individuals born in Sweden since 1932 and registered as living in Sweden since 1961 to their biological parents (38); 5) Prescribed Drug Register provides information on dispensed drugs to the entire Swedish population since July 2005. Active ingredients of drugs are coded according to the anatomical therapeutic chemical (ATC) classification system (39); 6) Longitudinal integration database for health insurance and labor market studies (LISA) was established in 1990 and contains annually updated data on highest level of education, civil status, unemployment, social benefits, and income for all Swedish residents aged 16 years or older (40); 7) National School Register (NSR) provides individual level data on final grades from school leaving certificates and eligibility to upper secondary school (41); 8) National Crime Register covers violent and non-violent crime convictions since 1973 (42).

The following were used as predictors. 1) Parental information: age and (maternal) weight at child birth, educational and marital status, criminal records, medical records; 2) family size and the numbers of siblings, full- and half-sibling medical and criminal records; 3) Family SES: neighborhood deprivation scores (NDEP, (43), family location (metropolitan area or not), family income, and social allowance received; 4) Index child information: perinatal records (child birth complications, APGAR scores, body measurements at birth), medical and criminal records. Medical records included inpatient and outpatient discharge records for 34 disorders and seven categories of prescription drugs. These disorders and prescription records were chosen based on prior knowledge of their relevance to ADHD or SUD. The complete ICD and prescription codes are listed in Supplementary file 1.

Missing information is common in register databases. All subjects in our cohort had one or more missing data point; less than 30% of the variables in our cohort were complete. Instead of removing subjects and variables with missing data, which could lead to a biased and reduced sample size and losing information, we handled missing data as follows. For categorical variables, we added a category to indicate “missing” status. Continuous variables with missing observations, such as body weight and family income, were first re-coded as categorical variables by quantiles splitting at 95, 75, 25 and 5% boundaries with a separate categorical code used for missing values. Continuous variables such as family income were recoded as categorical variables at each year to avoid the impact of inflation and changes in socioeconomic growth. Head circumference and birth weight were recoded as categorical variables at each gestational age (weeks). All of these transformed variables were one-hot encoded as dummy variables. The yearly index, neighborhood deprivation scores (NDEP) often had several years with missing records. Therefore, we summarized NDEP as mean, minimum and maximum values for three life-periods for the index child: perinatal (age 0-2), childhood (2-12), and adolescence/teenage (12-17). Diagnostic, prescription and criminal records were summed as the total numbers of records per year for the longitudinal model or summed for the above three periods for the cross-sectional model.

### Machine Learning Models

The sample was randomly divided into training (70%), validation (15%) and test subsets (15%). The training and validation sets were used to optimize the model parameters (which are estimated when the model is trained) and hyperparameters (which are set before the model is trained). The hyperparameters control structural features of the model such as the number of layers in a neural network or the number of trees in a random forests model. The test set was reserved only for evaluating the performance of the final models. Continuous variables were scaled between 0 and 1 based on minimum and maximum values of the training set.

We designed two types of prediction models: 1) a cross-sectional model to predict SUD diagnosis during age 18-19 and 2) a longitudinal, recurrent neural network (RNN) model to predict new SUD cases each year. For the cross-sectional model, we used Scikit-learn’s grid search algorithm to search the hyperparameter spaces for the random forest classifier (RF, (44) using the training and validation samples. We used the HyperOpt algorithm (45) to optimized the hyperparameters for the deep neural network model, multilayer perceptron (MLP). For the longitudinal model, we implemented the Long Short Term Memory (LSTM) network (46) to learn the corresponding input features at each year. HyperOpt was used to select the number of layers and number of LSTM units. For both the MLP and LSTM networks, we implemented the adaptive moment estimation (Adam) optimizer and used binary cross entropy as the loss function for HyperOpt tuning. Best models were chosen based on the lowest total validation loss and tested on the test set. All machine learning algorithms were written in Python 3.5.

### Diagnostic Accuracy Statistics

For the test set performance, we computed receiver operating characteristic (ROC) curves and used the area under the ROC curve (AUC) as our measure of accuracy. The AUC and its confidence intervals were calculated in Stata 14 using the empirical method and compared with nonparametric approach by DeLong et al (47). For the RF model, the final output from the test samples was the predicted mean probability of having an SUD diagnosis across all trees in the forest. For the MLP and RNN model, a sigmoid function implemented in the output layer generated a continuous score indicating the probability of the given participant having a SUD diagnosis in the predicted future time window. The distributions of the SUD probability scores for SUD and non-SUD subjects were compared using the two-sample Kolmogorov–Smirnov test. We used these probability scores to evaluate our models’ sensitivity, specificity, positive predictive power (PPP) and negative predictive power (NPP) in STATA 14.

### Learning curves

We used learning curve analysis to evaluate the model’s bias and variance (48). The learning curve plots the training and test set AUCs from different fractions of the training sample up to the total sample size. With increasing sample size, the training and validation scores should gradually converge. Ideally, the training and validation scores converge at a high level of accuracy, indicating that the model learns well from the training set and generalizes well to the test set. If the training and validation scores converge at a lower point, then the model has been underfit, i.e., it does not learn sufficiently from the training samples, but it can generalize this low level of fit to the test samples. If the training scores are high, validation scores are low, and the two scores don’t converge, then the model is overfitting the data and generalizes poorly. Inspection of the learning curves provides clues to how models might be improved in the future.

### Feature Importance Scores

We computed each feature’s importance score for the RF model. The feature importance score is the fraction of the sample’s predictions due to the feature. These scores add up to a total of 100% for each model. For individual features, the higher the value, the more important their contribution to the prediction model. We further evaluated the contribution of the top important features by fitting the models with or without these features.

## Results

### Cross-sectional model

The RF cross-sectional model achieved an AUC of 0.73 (95%CI 0.70-0.76) on the test samples when predicting SUD diagnoses at all visits during age 18-19 from prior data (Figure 1). For records having more than one SUD diagnosis, SUD diagnoses at prior visits are used as predictive features. The MLP model yielded a significant AUC that was nearly the same as the AUC for the RF model (0.71, 95%CI 0.68-0.74, Supplementary Figure 1). We, therefore, report the RF models for subsequent analyses.

**Figure 1.**
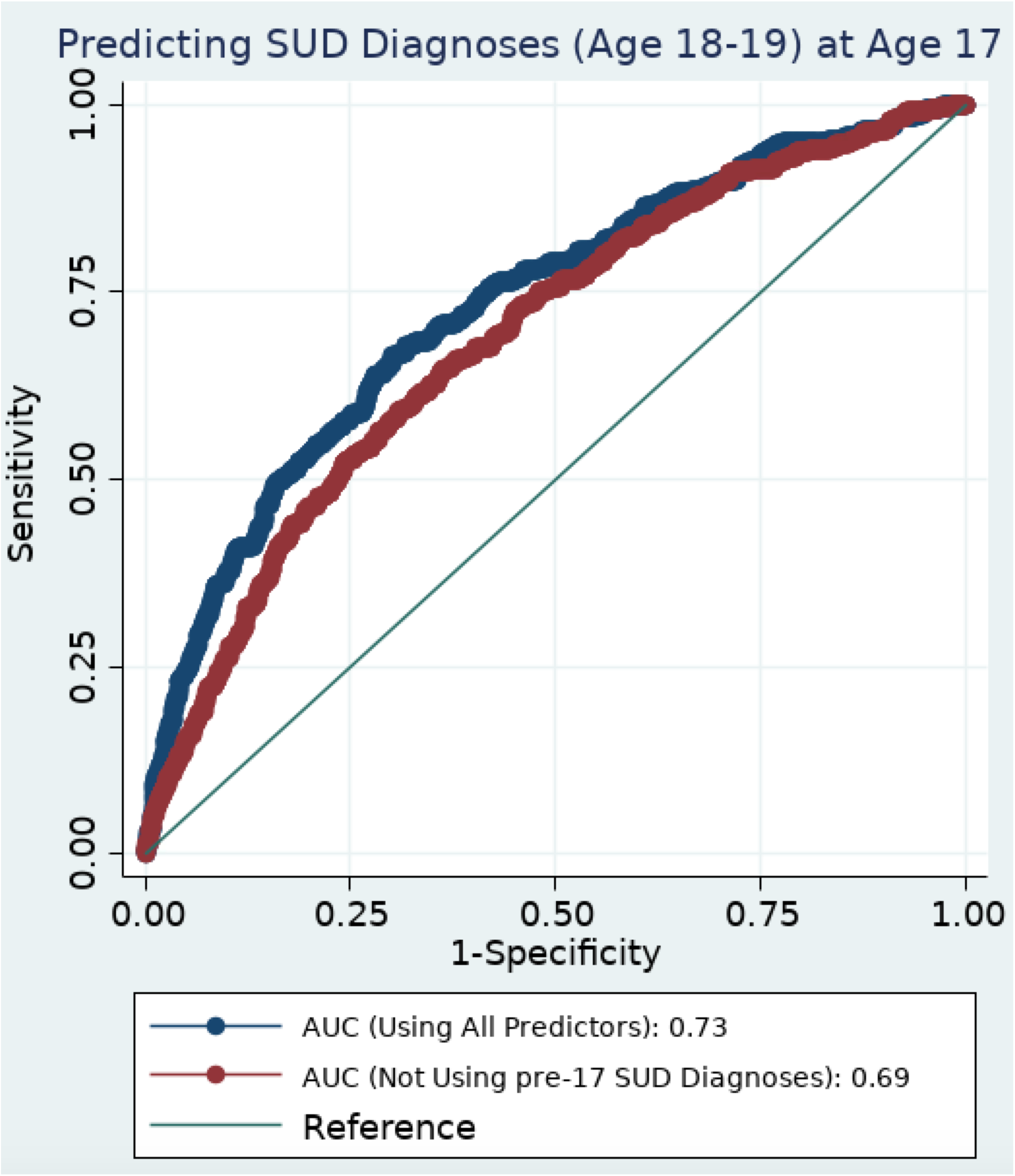
RF cross-sectional model prediction of all SUD diagnoses during age 18-19. Receiver Operation Characteristic (ROC) curves for the RF model were shown with or without using prior diagnosis of SUD as a predictor.

Having had a prior SUD diagnosis before age 17 was the most important predictor, accounting for 25% of predictive accuracy (Table 1). Figure 2, Left shows the feature importance scores across main categories. When prior SUD diagnoses for the index child were not used as a predictive feature, the remaining family members’ SUD diagnoses together accounted for 3% of predictive accuracy and other categories such as criminal records and family SES increased their contribution. When prior SUD diagnoses were removed from the prediction model, the AUC decreased to 0.69 (95%CI 0.66-0.72, Figure 1).

**Figure 2.**
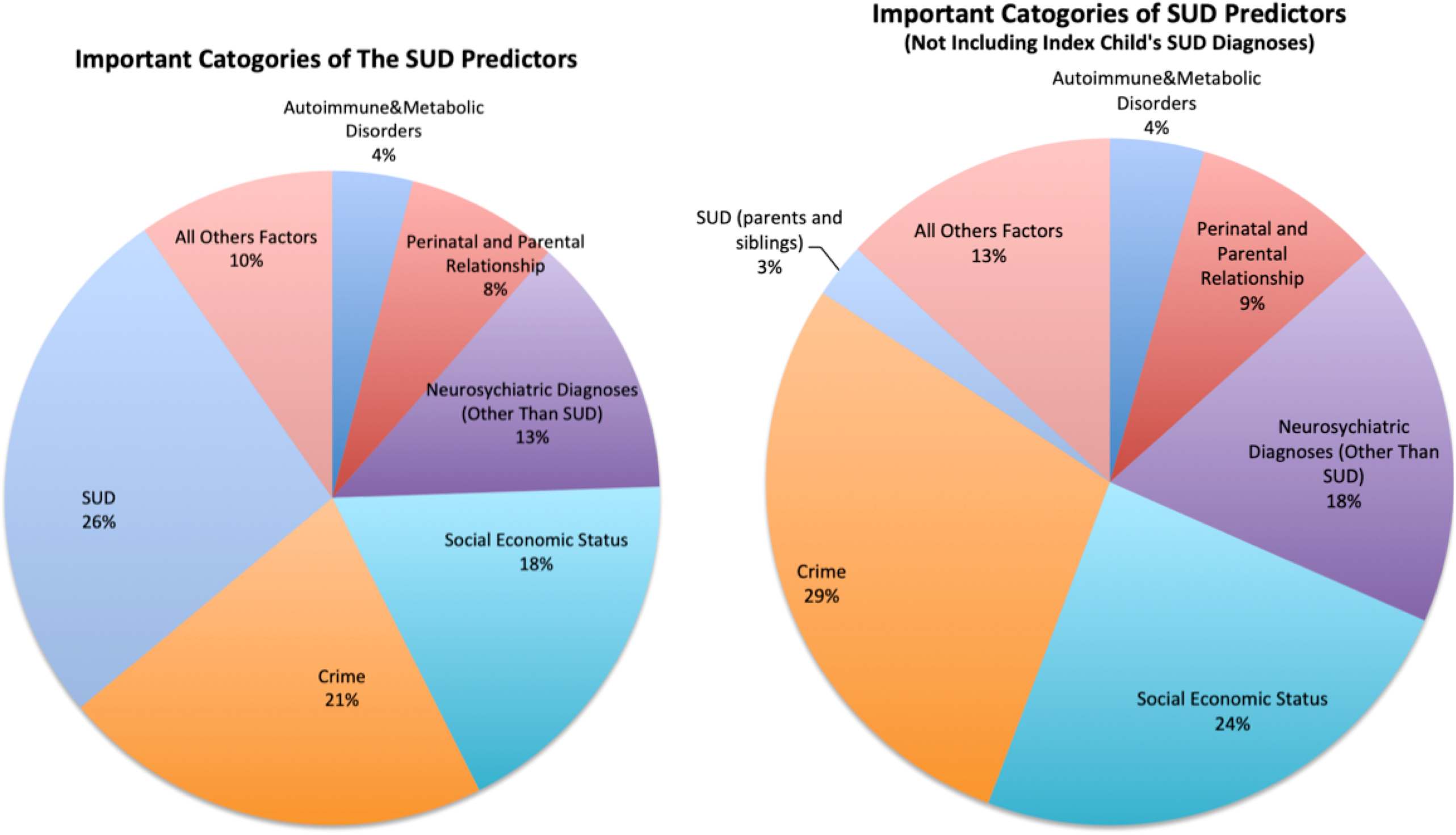
Feature Categories. Features important scores were combined into 7 main categories and their total contribution to the model predictions were plotted for the RF models with (Left) and without (Right) using prior diagnosis as a predictor.

When predicting only new onsets of SUD during age 18-19, the RF AUC was 0.67 (95%CI: 0.64, 0.71, Figure 3A). The sensitivity and PPP are plotted in Figure 3B with various probability cutoffs. Two examples of probability cutoffs were listed in Figure 3C with sensitivities 2.71% or 27.2% and their corresponding PPP at 54.6% or 20.4%.

**Figure 3.**
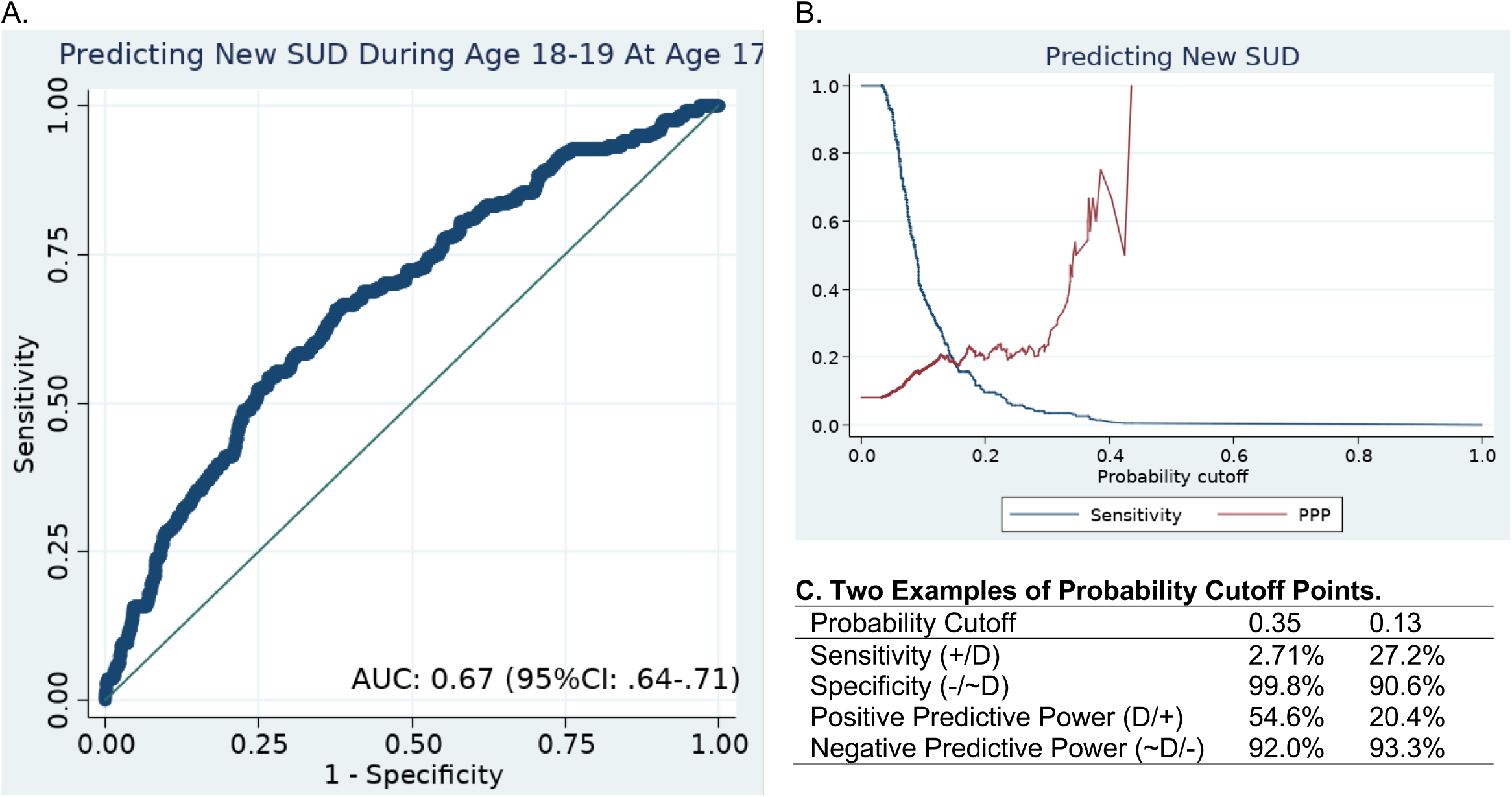
RF cross-sectional model predicting new SUD cases during age 18-19. **A**. Receiver Operation Characteristic (ROC) curve. **B**. Sensitivity and PPP plotted over various of probability cutoff points. **C**. Sensitivities, Specificities, PPPs and NPPs at two example probability cutoffs. Note 1: D, SUD-positive individuals; ∼D, SUD-negative individuals; +, positive predictions; -, negative predictions. Note 2: The sensitivity is the percentage of SUD diagnoses predicted among true SUD subjects. The specificity is the percentage of negative SUD diagnoses predicted for true non-SUD cases. The PPP is the percentage of true SUD in among the predicted SUD diagnoses. The NPP is the percentage of true non-SUD cases among those predicted not to have SUD diagnoses.

### Top Features

Table 1 lists the top 20 most important features. Besides a prior SUD diagnosis, the most important features were teenage criminal records (from onset age 15 up to age 17) and a childhood (age 2-12) ADHD diagnosis, followed by stimulant treatment prior to age 18, diagnosis of anxiety disorder and SES indices (such as family income and neighborhood deprivation scores) during teenage years (age 12-17). Father’s non-violent crimes before birth and SES indices, as well as ADHD diagnosis during teenage years were also among the top 20 list but ranked lower. When we removed the top 10 most important features from the model, prediction accuracy for new SUD cases was significantly reduced (AUC 0.59, 95%CI: 0.56-0.63, χ^2(1)^ =16.2, p=0.0001), although the prediction was still significantly above the chance AUC of 0.5 (Supplementary Figure 2). Excluding 10 more features did not significantly reduce further the AUC (0.58, 95%CI 0.54-0.62). The importance of the top 10 ranked features was further confirmed by the significance and magnitude of the AUC when using only these 10 features were in the model (0.66, 95%CI 0.62-0.70). We found similar results using the top 20 features (0.67, 95%CI: 0.64-0.71).

### Longitudinal model

Figure 4A illustrates the longitudinal model design depicting the input predictors at each age. The prediction AUCs at each age are shown in 4B, for predicting new SUD diagnoses one, two, five or ten years in the future. The average AUCs were similar (0.65∼0.66) for all the intervals and majority of the AUCs were significant with their 95% CI intervals above 0.5. We compared the two-year outlook prediction of new SUD cases at age 17 for the cross-sectional and longitudinal models. Supplementary Figure 3 shows that both models have significant AUCs that were above 0.5. However, the cross-sectional model had a significantly higher AUC than the longitudinal model (χ^2(1)^ =6.60, p=0.01).

**Figure 4.**
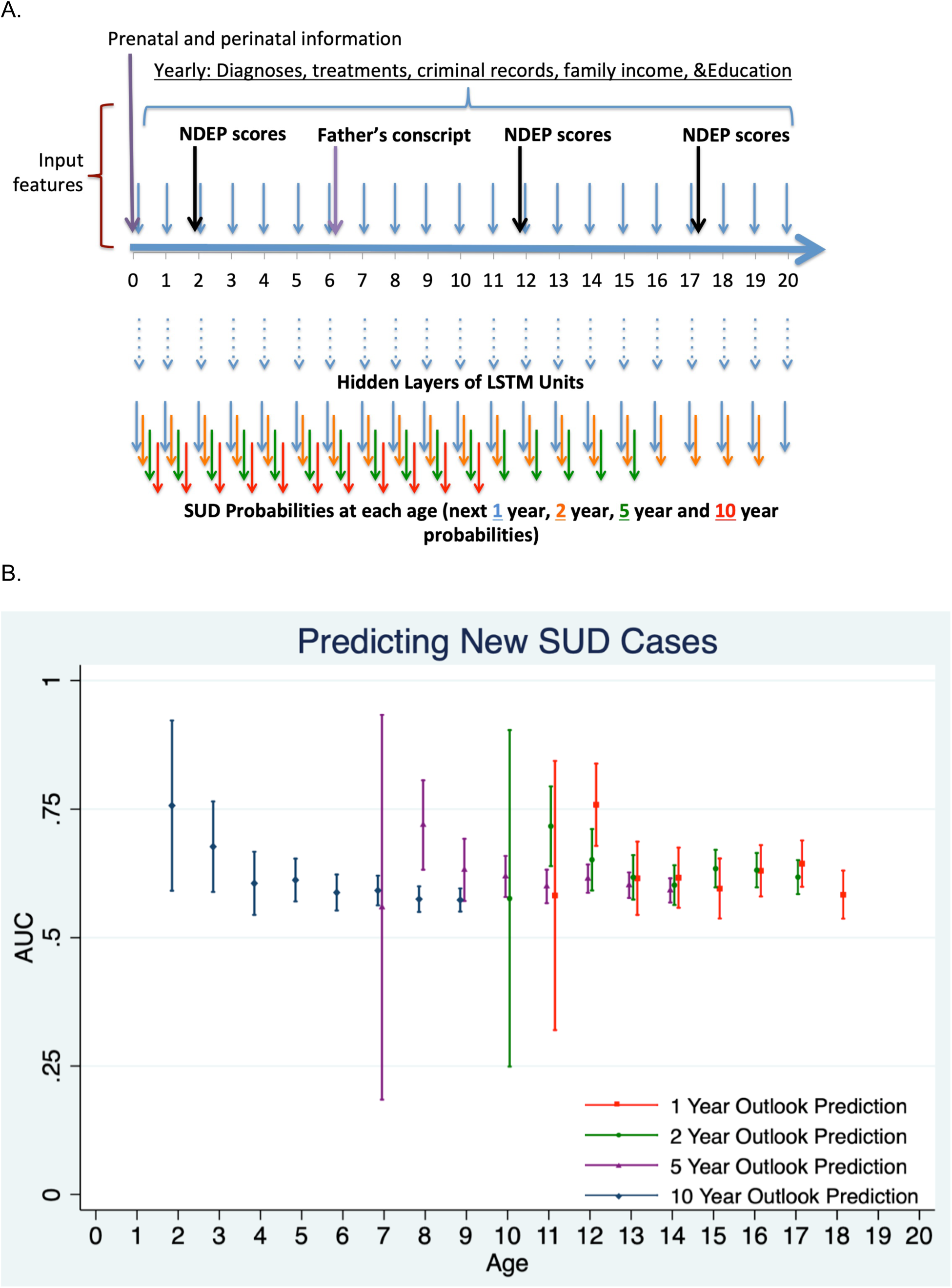
Longitudinal model predicting new SUD diagnoses at each age. **A**. Model architecture. **B**. AUC at each age for 1, 2, 5 and 10 year outlook predictions.

### Learning curve analysis

The learning curve was plotted for the RF cross-sectional model (Supplementary Figure 4). Training and validation AUCs gradually converged with the increased samples used. However, their final scores did not fully converge. In addition, the validation AUCs plateaued rather quickly. The learning curve characteristics suggest that more training samples and additional informative features are both needed to improve the prediction accuracy and reduce over-fitting.

## Discussion

Using Swedish population register data, we used two different models to predict SUDs in ADHD youth. The cross-sectional model significantly predicted the probability of having SUD during ages 18-19. The longitudinal model predicted short- and long-term risks for future new diagnoses at each age. Both models yielded significant predictions. Notably, the longitudinal model was able to predict future SUD diagnoses at young ages, many years prior to their ages at first SUD diagnosis.

This study is the first to apply machine learning algorithms to predict a serious and public health relevant outcome in the context of ADHD. We evaluated the potential clinical utility of the prediction models by computing sensitivity, specificity, PPP and NPP at several cutoff points (Figure 3B and C). Ideally, a prediction model would identify most patients who would go on to future substance use (high sensitivity) and few who would not (high PPP). The sensitivities, specificities, PPPs and NPPs we reported are not ideal. They do, however, indicate that large-scale data combined with machine learning may eventually arrive at clinically useful prediction models. Analyses of the full electronic health records ascertained from the actual health care system or even more detailed predictor information from linked population-register are two strategies to improve the predictive power.

Future research also needs to carefully consider clinically useful cutoff points for the obtained risk scores. For example, Figure 3B shows that using a cutoff point with a sensitivity of 2.7%, defines a sample in which 54.6% (the PPP at that cutoff point) will have a subsequent SUD diagnosis. If we instead use a cutoff point with a sensitivity of 27.2%, the PPP decreases to 20.4%, which means that only 1 in 5 patients defined by the model as being at risk for SUD are truly at-risk. Although this cutoff point gives a low PPP, it could be useful because the burden of data collection is low (the data are already available in the medical record) and the results can be used for economical and non-invasive interventions such as parent and patient education or more frequent monitoring of high-risk patients.

The feature importance scores provide insights into which features provided the most information about the future risks of SUDs. They are, however, not adjusted for confounders and should be interpreted as useful predictors but not, based on our analyses, etiologic risk factors. Furthermore, although the feature importance scores from the Random Forests algorithm will extract important features, we cannot conclude that features not extracted are not relevant to SUD. For example, if two features are highly correlated, the one which is most predictive will be deemed important. The other one will have a low importance score because it is not useful after the first one has been selected into the model (49). Therefore, although the importance scores are useful for explaining the performance of our models, they should not be used to make relative comparisons between features with regards the degree of risk they impart for SUD. In other words, the top features may mask other smaller but important predictors of SUDs. Because of such correlations, when we dropped the prior diagnoses of SUD from our models, their AUCs did not drop dramatically, indicating a substantial amount of redundant information in the remaining feature set.

Consistent with prior work (50, 51) (52), we found that prior committed crimes (both non-violent and violent crimes) during teenage years contributed 16% to the predictive accuracy. Indeed, in our sample, those who committed crimes during teenage years had three times higher SUD prevalence (29.4%) during age 18-19 than those who did not have criminal history (prevalence 9.6%). Many previous studies have reported that low family socioeconomic status is associated with SUDs (e.g., 15, 53). Indeed, numerous SES features in our study, including family income and neighborhood deprivation scores, had high feature importance scores. Altogether, features from the SES category accounted for 18 to 24% of predictive accuracy (Figure 2B). Having had an ADHD diagnosis in childhood was ranked fourth in feature importance. Interestingly, children who had been diagnosed with ADHD during childhood had lower risks for SUD (5.2% would have SUD during 18-19) than those who were diagnosed with ADHD during adolescence (9.4%). Those who were diagnosed with ADHD during 18-19 had the highest SUD prevalence 14.3%. This could reflect changes in diagnostic and treatment practices with calendar time but it is possible that delayed diagnoses of ADHD increase the risk for SUD due to delayed treatment. This idea is supported by many studies showing that the treatment of ADHD in youth leads to lower risks for outcomes such as criminality (52), traffic accidents (54), smoking (26) and SUDs (27, 28).

Our work has some limitations. First, our learning curve analyses suggest that improvements in predictive accuracy will require additional features and a larger sample size. It is also possible that major improvements to prediction will require a different source of data, such as biomarker or behavioral assays, which are not available in the registries. Secondly, we have only information available up to age 20 for all patients. Therefore, we are only predicting the SUD onset risks up to age 20 and can’t draw conclusions about predictive accuracy for older ages. Thirdly, our sample only contains patients with a diagnosis of ADHD. We do not know if our model predictions would generalize to the population. Future studies to predict the SUD in the entire population would be useful. Finally, our feature importance analysis, albeit informative for prediction mechanisms, does not provide direct evidence for etiological risk factors for SUD. However, future research based on new ideas derived from our feature importance analysis could provide evidence clarifying if there are any causal mechanisms and perhaps discover novel links.

Despite these limitations, our results suggest that population registry data are useful for machine learning algorithms to predict the future onset of SUDs when the actions taken based on the predictions are neither costly nor invasive. Future work should focus on improving the sensitivity and positive predictive value by including more detailed information from predictors.

## Acknowledgements

This project has received funding from the European Union’s Horizon 2020 research and innovation programme grant agreement No 667302. This publication reflects only the author’s view and the European Commission is not responsible for any use that may be made of the information it contains. Henrik Larsson acknowledge financial support from the Swedish Research Council (2018-02599) and the Swedish Brain Foundation (FO2018-0273). Additional support was provided by Shire Development, Inc. We thank Machine2Learn for programming support.

## Disclosure

Yanli Zhang-James, Qi Chen, Ralf Kuja-Halkola and Paul Lichtenstein declare no conflict of interest. In the past year, Dr. Faraone received income, potential income, travel expenses continuing education support and/or research support from Vallon, Tris, Otsuka, Arbor, Ironshore, Shire, Akili Interactive Labs, VAYA, Ironshore, Sunovion, Supernus and Genomind. With his institution, he has US patent US20130217707 A1 for the use of sodium-hydrogen exchange inhibitors in the treatment of ADHD. Dr. Larsson has served as a speaker for Evolan Pharma and Shire and has received research grants from Shire; all outside the submitted work.

## List of Tables

**Supplementary File 1** Complete list of ICD codes for diagnosis and prescription codes used.

---

**Supplementary Table S1.** Prescription medications for SUD treatment.

| Medications |  | ATC code |
| --- | --- | --- |
| Medications for treatment of SUD |  | N07B |
| Medications for treatment of alcoholism |  |  |
|  | acamprosate | N07BB03 |
|  | naltrexone | N07BB04 |
|  | disulfiram | N07BB01 |
| Medications for treatment of drug abuse |  |  |
|  | buprenorphine | N02AE01 |
|  | buprenorphine+naltrexone | N07BC51 |
|  | methadone | N07BC02 |

**Supplementary Table S2.** Numbers of SUD diagnoses prior to age 17 and during age 18-19.

| SUD Diagnosis<br>Prior to 17 | No SUD diagnosis<br>(age 18-19) | Has SUD diagnosis<br>(age 18-19) | Total |
| --- | --- | --- | --- |
| 0 | 16,133<br>(84.1%) | 1,134<br>(5.9%) | 17,267<br>(90%) |
| 1 | 1,358<br>(7.1%) | 559<br>(2.9%) | 1,917<br>(10%) |
| Total | 17,491<br>(91.2%) | 1,693<br>(8.8%) | 19,184<br>(100%) |

**Supplementary Figure S1.**
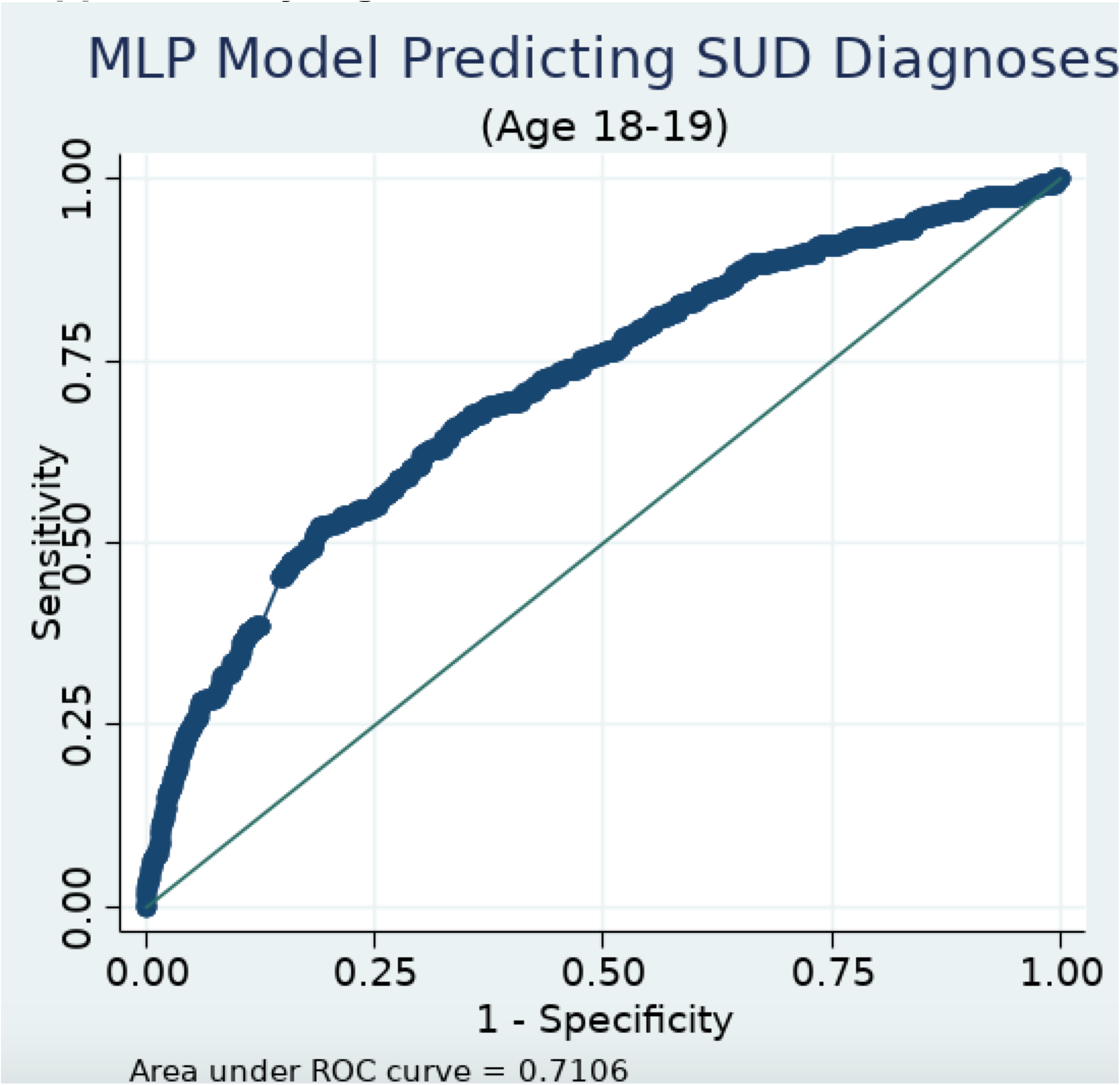
Receiver Operation Characteristic (ROC) curve for MLP cross-sectional model at age 17.

**Supplementary Figure S2.**
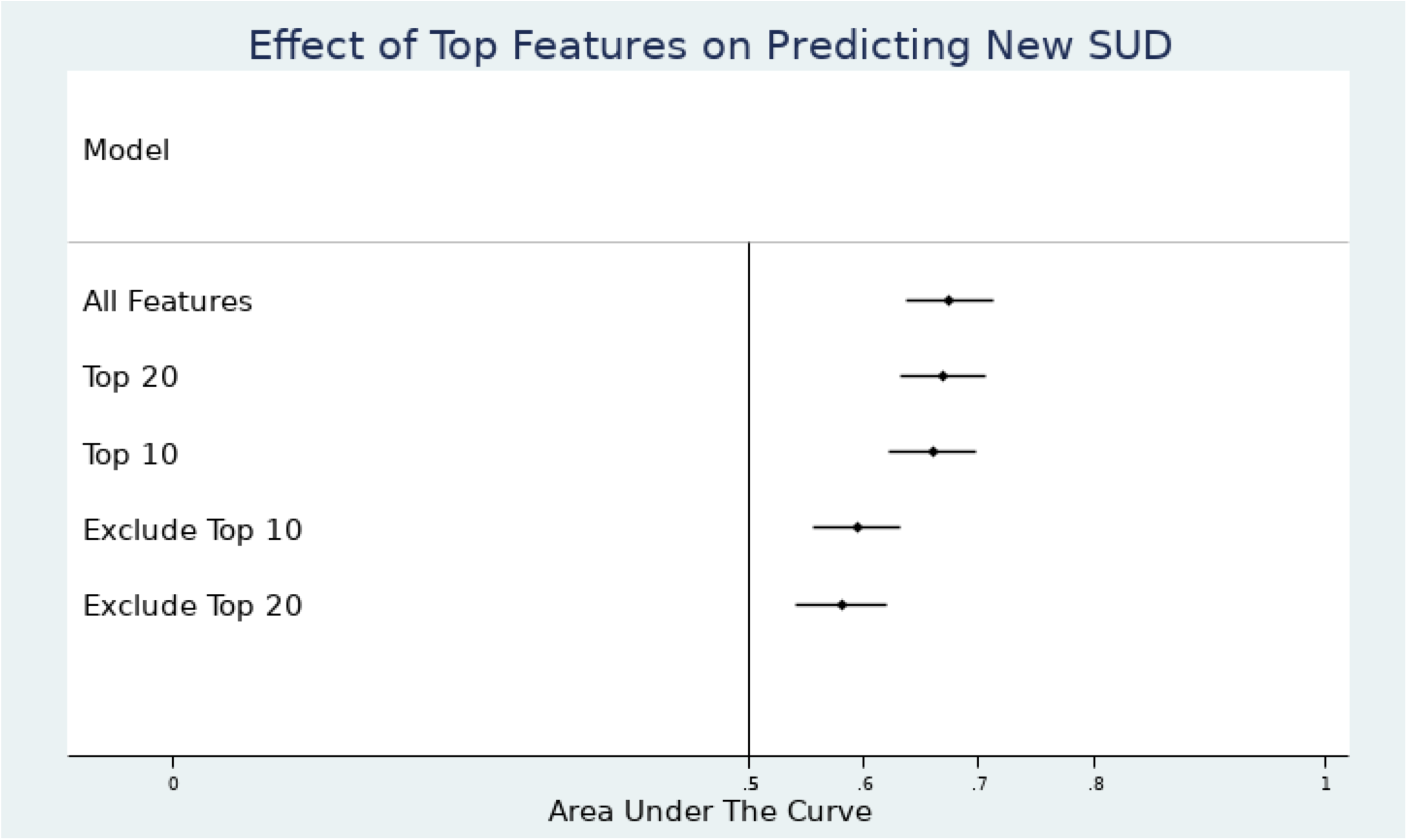
Effects of Top Features on Prediction of New SUD Cases. AUC and 95%CI were plotted for predictions that used all the features, only the top 10 or 20 features, or all the features except with the top 10 or 20 removed.

**Supplementary Figure S3.**
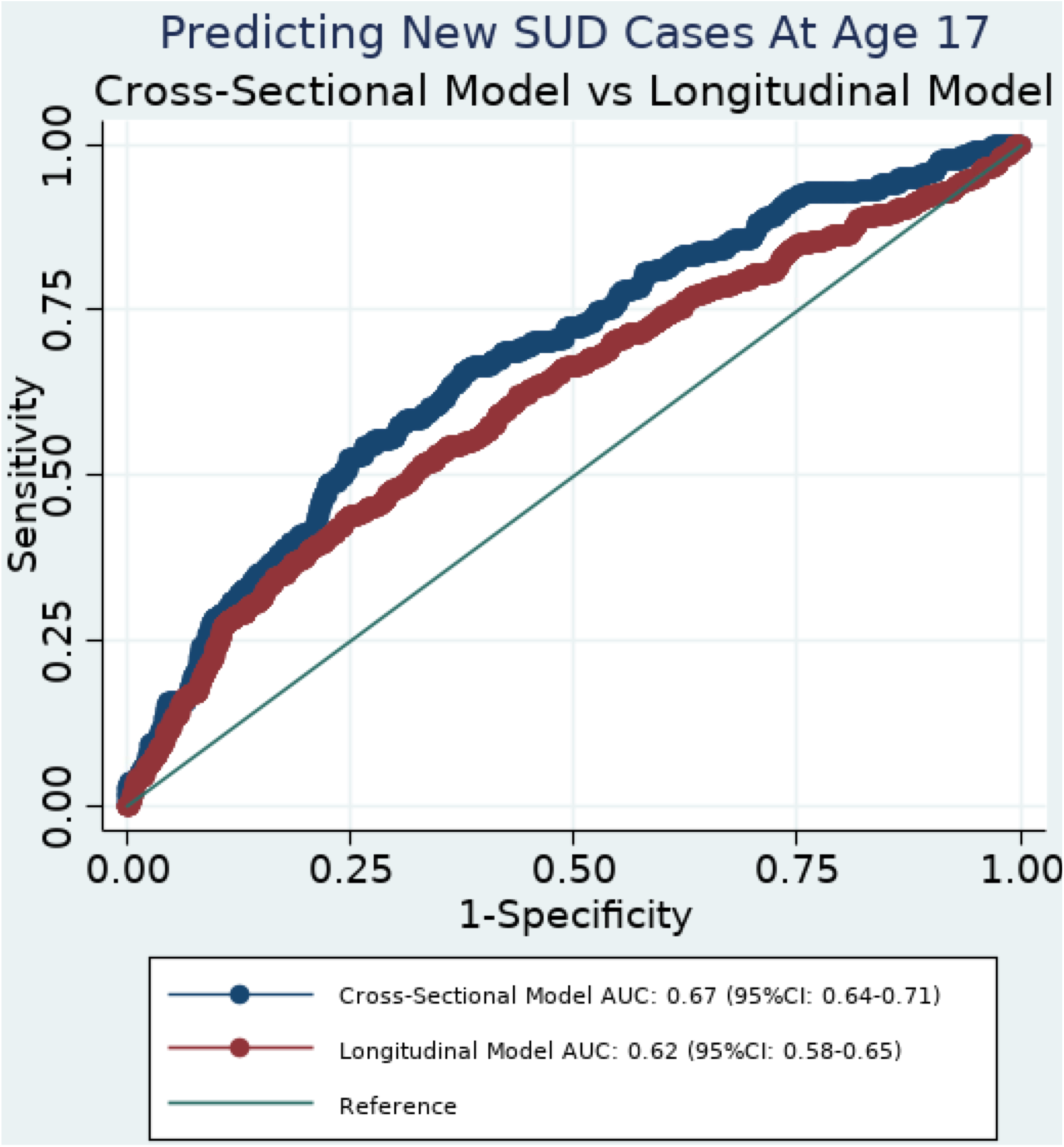
Predicting new SUD cases at age 17: cross-sectional model vs longitudinal model. Cross-sectional model AUC was significantly higher than that of the longitudinal model (Chi^2(1)^ =5.17, p= 0.023). However, both AUCs were significantly above 0.5, where prediction is by chance.

**Supplementary Figure S4.**
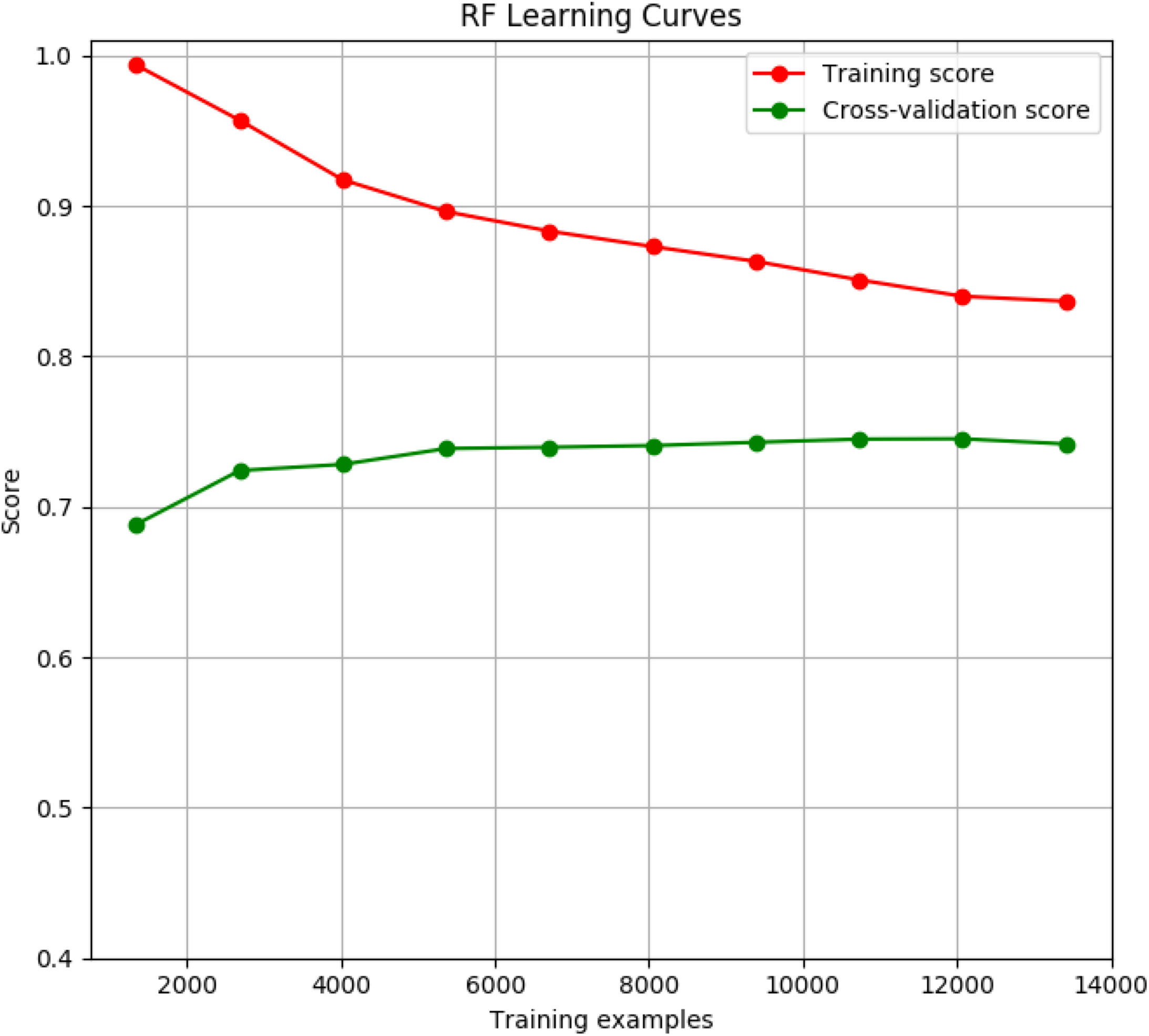
**Learning curves** were plotted for the RF cross-sectional model. X axis indicates the numbers of samples from the total training samples were used in each itineration and y axis were the training and validation AUCs.

